## Supplemental figures and legends for "Dual engagement of the nucleosomal acidic patches is essential for deposition of histone H2A.Z by SWR1C"

Baier et al., Supplemental Figure 1

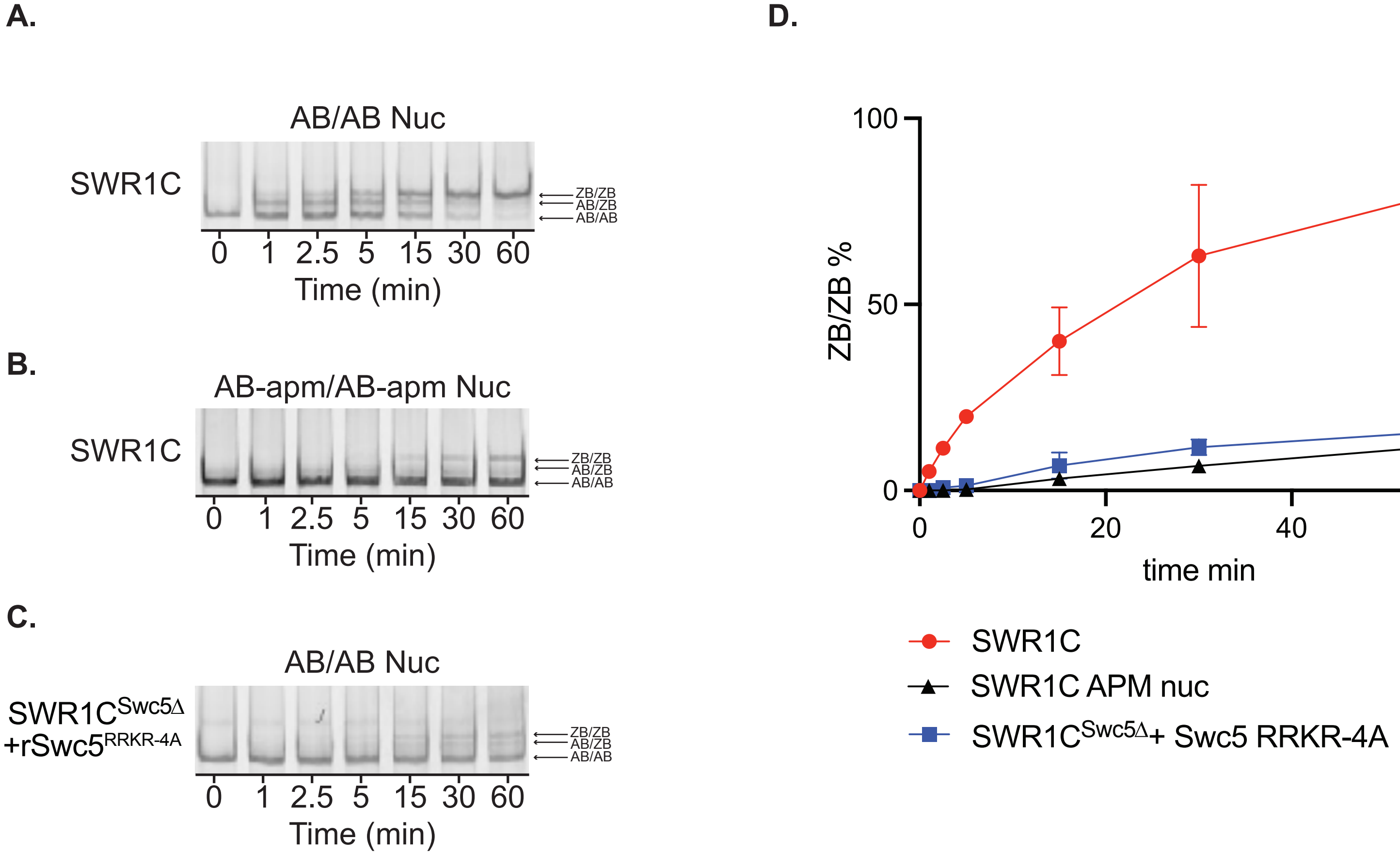

**Supplemental Figure 1**

**Gel-based assay for H2A.Z deposition (related to Figure 1).** SWR1C mediated ZB deposition was measured by incorporation of H2AZ$^{3x\text{-}FLAG}$/H2B dimers at various time points on 77N0 AB/AB **(A)** or 77N0 AB-apm/AB-apm **(B)** nucleosomes. SWR1C$^{Swc5\Delta}$ +rSwc5$^{RRKR\text{-}4A}$ mediated ZB deposition on 77N0 AB/AB nucleosomes was also measured **(C)**. Percent of remodeled nucleosome in each condition was quantified using ImageQuant and plotted in **(D)**.

Baier et al., Supplemental Figure 2

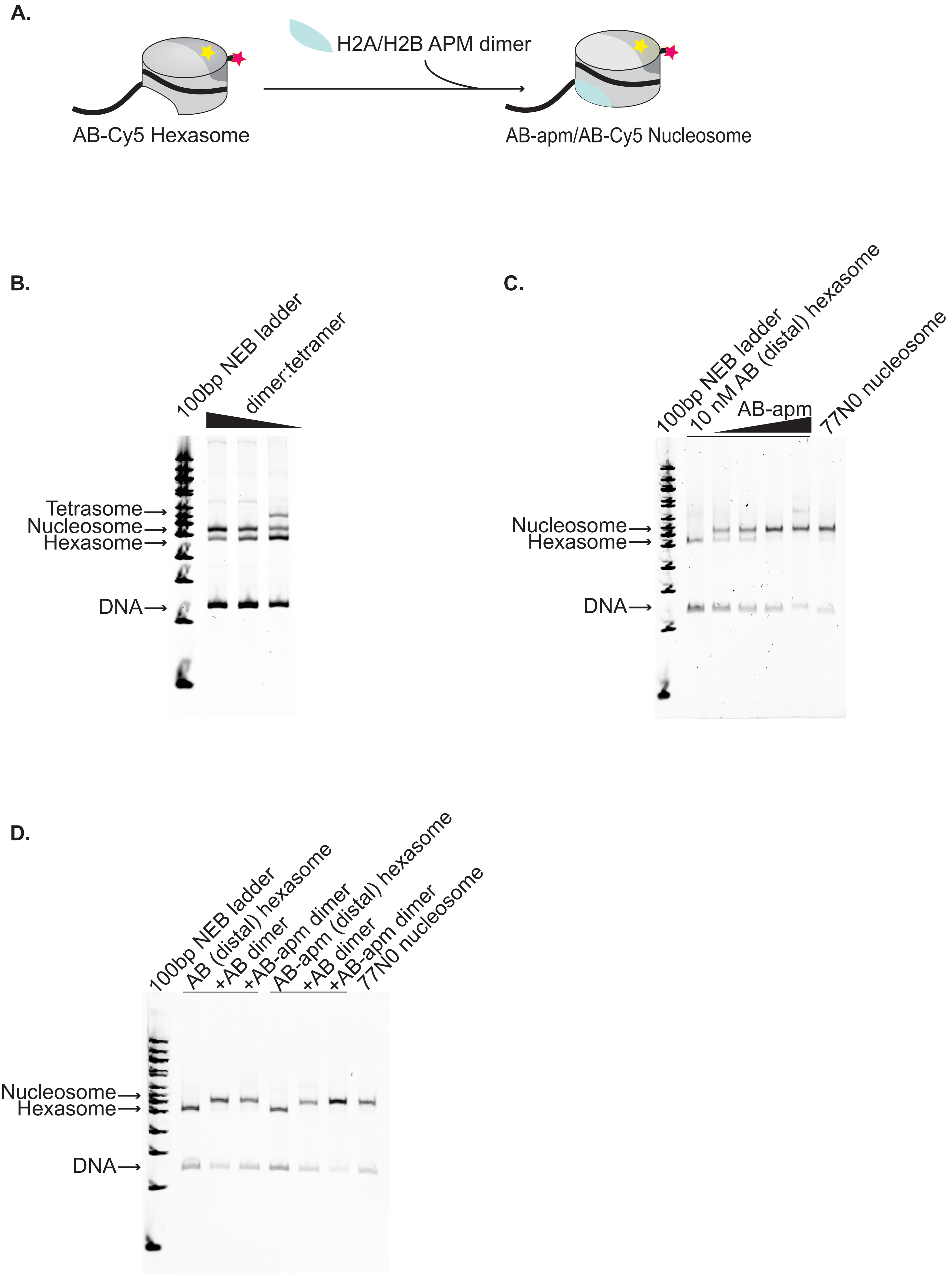

**Supplemental Figure 2**

**Strategy for assembly of asymmetric nucleosomes (A)** Schematic of assembly of asymmetric nucleosomes from purified hexasomes. Hexasomes (represented by partial cylinder, H3/H4 tetramer in light grey, AB dimer in dark grey, Cy5 yellow star, 77N0 DNA by thick black line, and Cy3 by pink star) spontaneously assemble into nucleosomes upon addition of free dimer (AB-apm, shown added in light blue). **(B-D)** give representative examples of the checkpoints performed during assembly of asymmetric nucleosomes. **(B)** For each hexasome needed for biochemical assays, the ratio of dimer to tetramer was varied during reconstitution to optimally bias the reaction toward hexasomes and assessed by native-PAGE with a tetramer:DNA ratio of 1.4 and a dimer:DNA ratio of 1.4, 1.8, and 2.4. **(C)** After purification of hexasomes, contralateral dimer to be placed was titrated to determine optimal conversion to nucleosome and assessed by native-PAGE with a canonically reconstituted 77N0 or 0N0 nucleosome as appropriate added for comparison with a ratio of dimer to hexasome of 0.9, 1, 1.1, and 1.2. **(D)** As a final check before performing biochemical assays, each hexasome was run on a native-PAGE gel with nucleosomes formed by dimer addition and a canonically assembled nucleosome for comparison.

Baier et al., Supplemental Figure 3

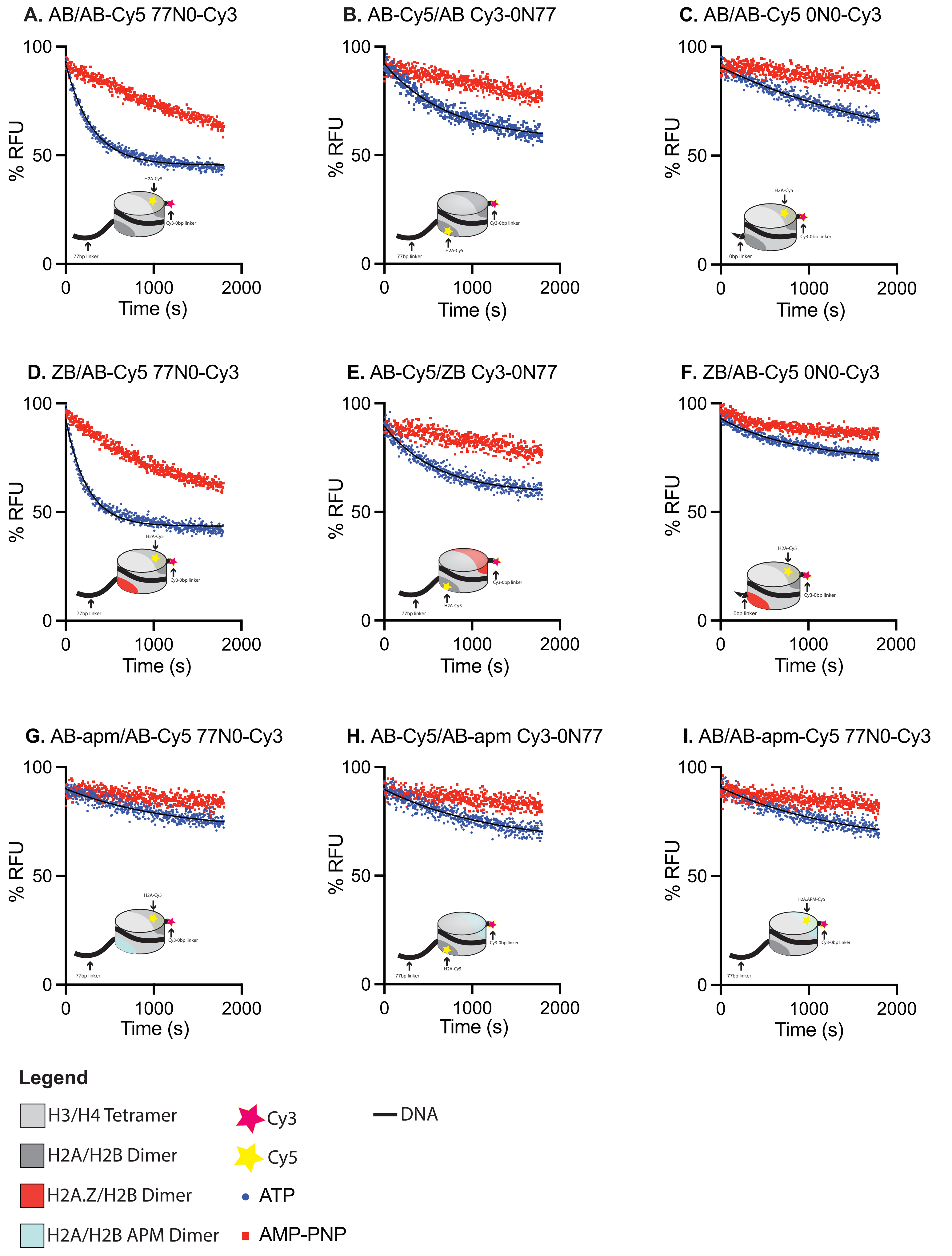

**Supplemental Figure 3**

**FRET-based dimer eviction (related to Figure 1) (A-I)** Graphs of dimer eviction data used to calculate rates in Figure 1C and 1D. Reactions were carried out under single turnover conditions (50 nM SWR1C, 10 nM nucleosomes, 60 nM ZB dimers) in the presence of ATP (blue) or AMP-PNP (red) for each of the asymmetric nucleosome (listed linker proximal/ linker distal / DNA template at top left of each graph) and loss of FRET was measured over time. **(A-F)** Rates for ATP reactions differed significantly from AMP-PNP while in **(G-I)** there was no significant difference. **A, D** and **F** have six replicates while the rest are done in triplicate. Note that the AMP-PNP reactions in panels A and D showed higher rates of FRET loss compared to all other substrates. This was reproducible in 3 different nucleosome preparations and in all replicates. This loss of FRET was not observed in the no enzyme control, indicating enzyme-dependent destabilization (see source data file). These higher 'background' rates are not observed when this same DNA template is reconstituted with Cy5-labelled octamers; thus, the apparent instability of these nucleosomes seems inherent to the hexosome assembly method.

Baier et al., Supplemental Figure 4

**A.**

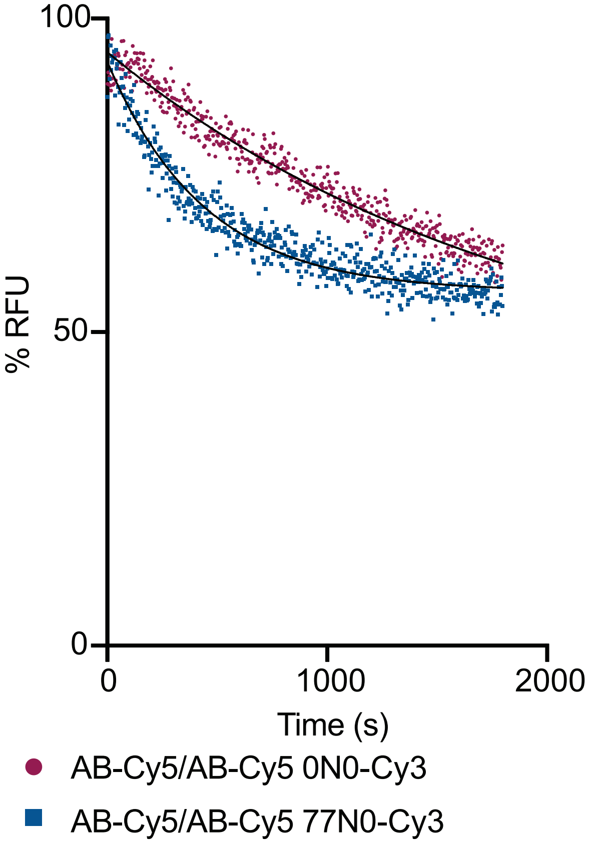


**B.**

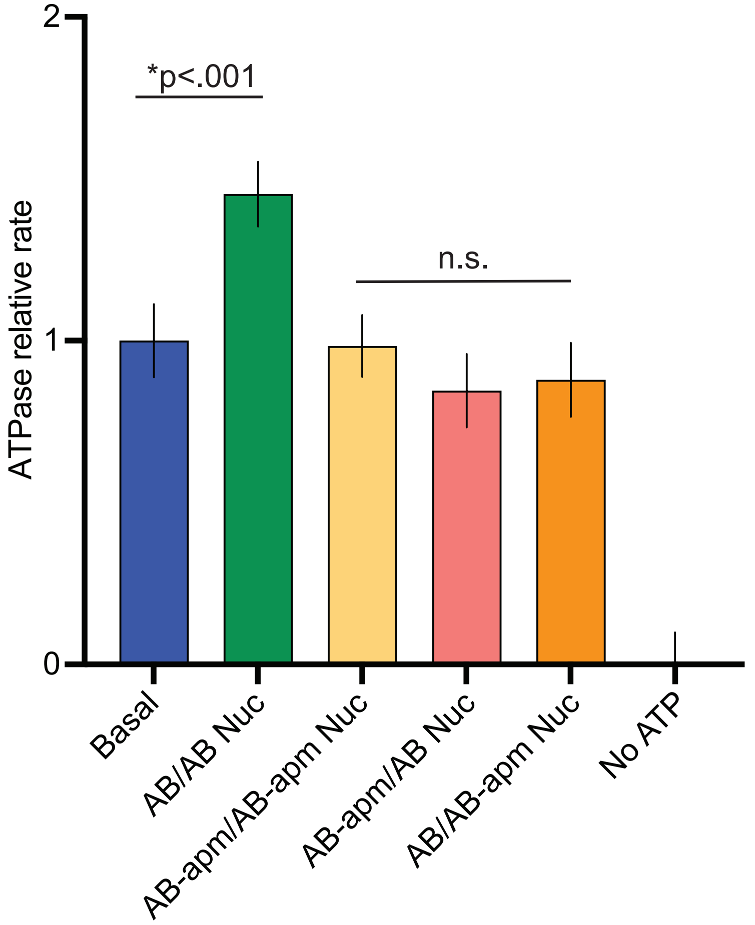


**C.**

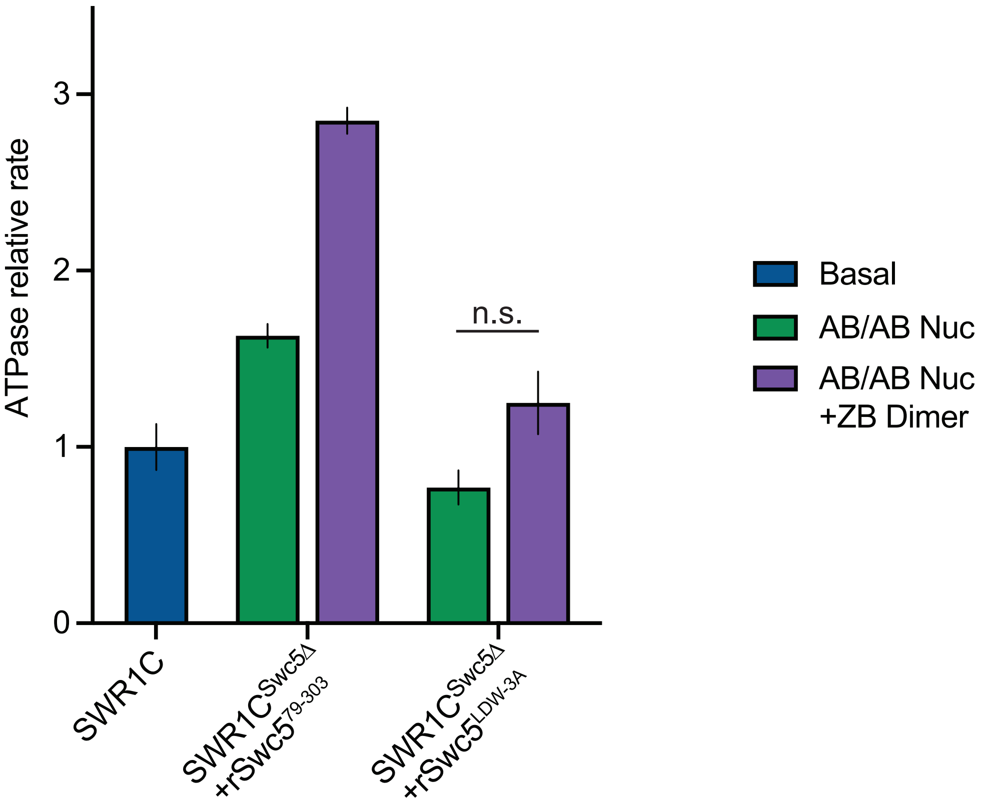

**Supplemental Figure 4**

**Dimer eviction and ATPase assays. (A)** Graph of dimer eviction data carried out at high enzyme concentrations (200 nM SWR1C, 10 nM nucleosomes, 250 nM ZB dimers, 1 mM ATP) on AB/AB-Cy5 0N0-Cy3 nucleosomes (maroon) or AB/AB-Cy5 77N0-Cy3 nucleosomes (blue). SWR1C evicted AB-Cy5 dimers significantly faster in the presence of linker DNA. **(B)** ATPase activity of SWR1C was measured by phosphate binding protein assay in the presence of various asymmetrically assembled 77N0 nucleosomes and normalized to basal activity. AB/AB nucleosomes (green bar) stimulated the ATPase rate as compared to basal SWR1C activity (blue bar). SWR1C activity in the presence of nucleosomes containing a linker proximal (yellow), distal (salmon) acidic patch mutant, or both (orange) acidic patches mutated did not differ significantly from basal SWR1C rates. **(C)** Nucleosomal stimulation of SWR1C ATPase activity with (purple bars) or without (green bars) ZB dimers added was measured for SWR1C, $SWR1C^{Swc5\Delta}$ +$rSwc5^{79\text{-}303}$, and $SWR1C^{Swc5\Delta}$ +$rSwc5^{LDW\text{-}3A}$ using a phosphate sensor assay. Calculated rates were normalized to basal SWR1C activity (blue bar). Stimulation of ATPase activity was restored to the Swc5Δ complex by the $Swc5^{79\text{-}303}$ derivative, but not by the $Swc5^{LDW\text{-}3A}$ derivative.

Baier et al., Supplemental Figure 5

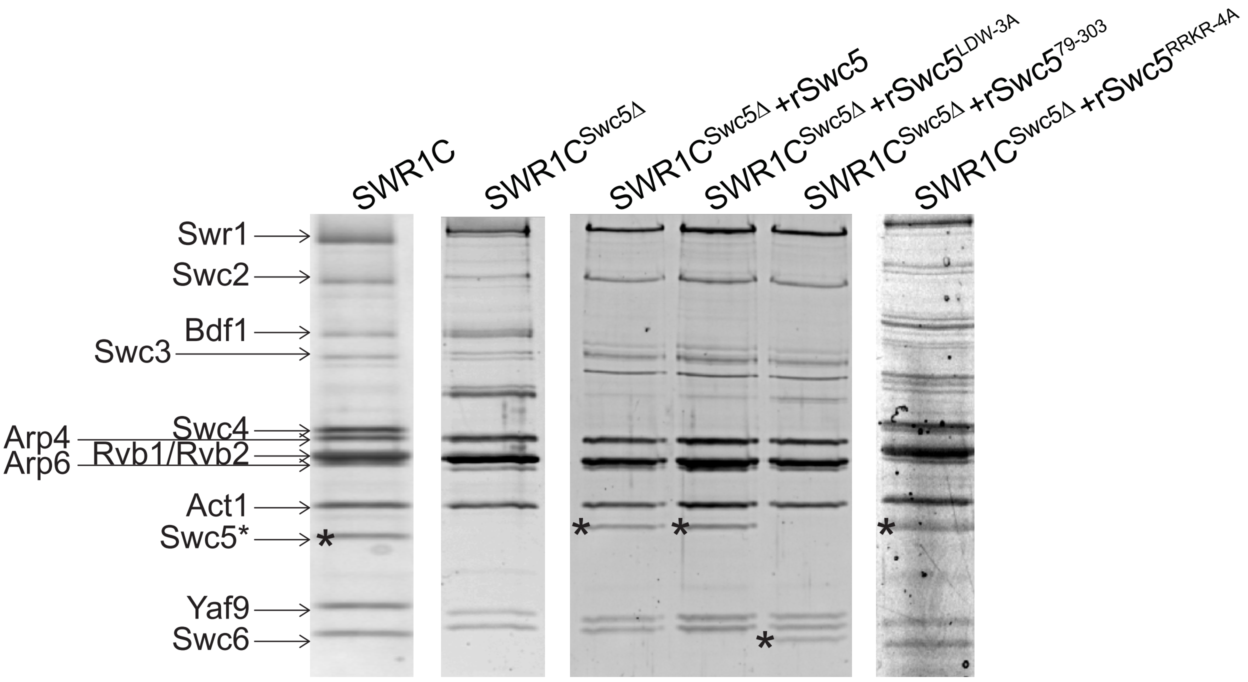

**Supplemental Figure 5.**

**Reconstitution of SWR1C with Swc5 derivatives. (A)** SDS-PAGE gel showing wild-type SWR1C, SWR1C$^{Swc5\Delta}$, and SWR1C$^{Swc5\Delta}$ with various recombinant Swc5 reincorporated (Swc5 denoted by asterisk).

Baier et al., Supplemental Figure 6

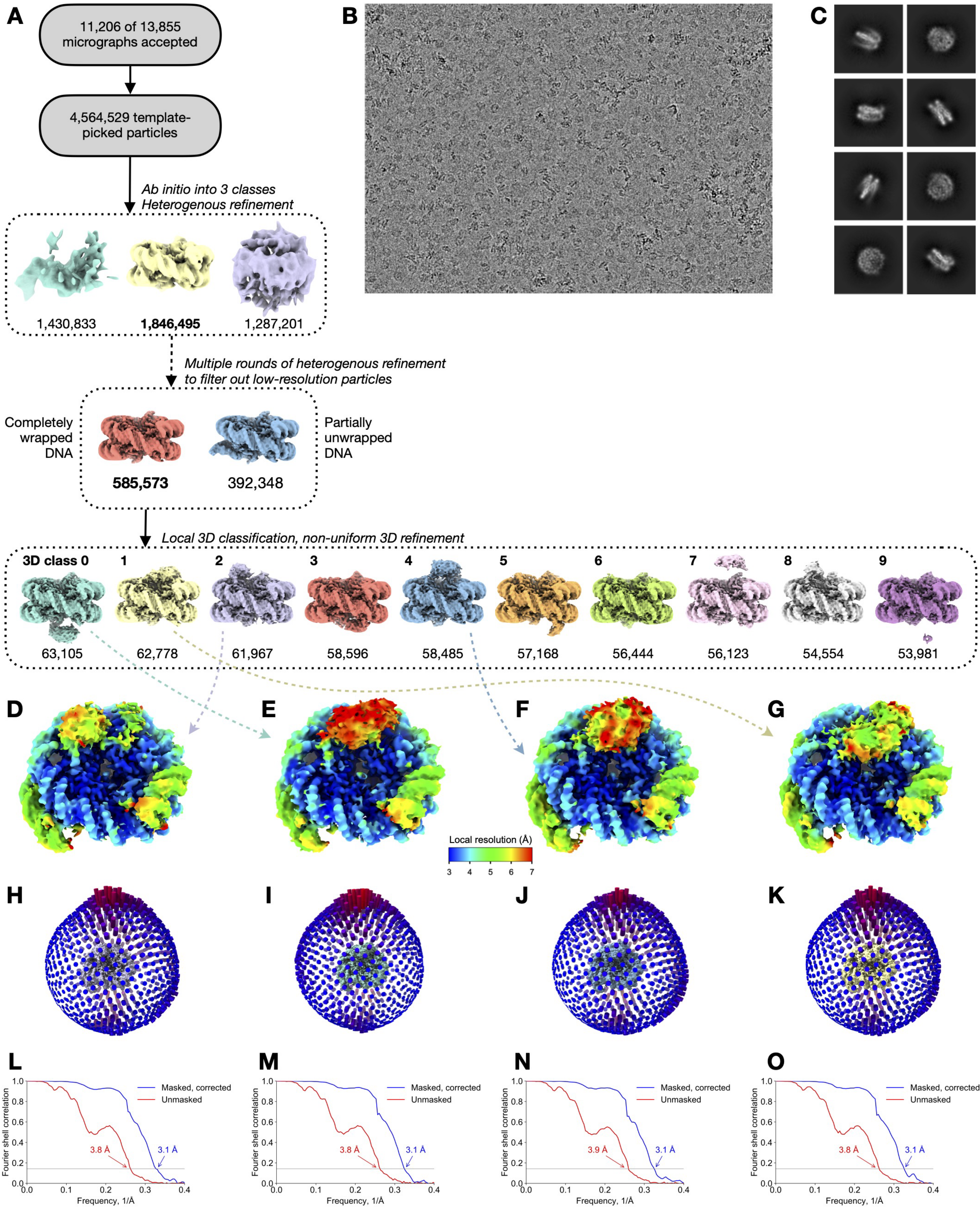

**Supplemental Figure 6.**

**Cryo-EM analysis of Swc5/nucleosome complex. (A)** Flowchart of data processing and particle classification. The 3D classes marked with letters E-G correspond to the four selected reconstructions in the panels below and in Figure 6. **(B)** Representative cryo-EM micrograph. **(C)** Selected class averages from 2D classification of the 1,846,495 nucleosomal particles obtained after the first round of heterogenous 3D refinement. **(D-G)** Selected 3D reconstructions (the same as in Figure 6) colored by local resolution. **(H-K)** Euler angle distributions of the particles used in the reconstructions in panels D-G. The view angle is the same as in panels D-G. **(L-O)** Fourier shell correlation plots of independently refined half maps corresponding to reconstructions in panels D-G. Resolution is determined at FSC=0.143.

**Supplemental Movie 1.**

**Conformational heterogeneity of Swc5.** The 3D class depicted in Figure 6C and Supplemental Figure 1F,J,N was subjected to cryoSPARC's 3D Variability Analysis. Variability was solved in 2 modes, filtered to 7 Å, and a white noise model was used. The animation displays the primary component of heterogeneity in the subset – Swc5 moiety swinging between H4 N-terminal tail on the left and H2A/B dimer on the right.

**601 positioning sequence**
GTGAAATACCGCACAGATGCGTAAGGAGAAAATACCGCATCAGGCGCCATTCGCC
ATTCAGGCTGCGCAACTGTTGGGAAGGGCGATCGGTGCGGGCCTCTTCGCTATTA
CGCCAGCTGGCGAAAGGGGGATGTGCTGCAAGGCGATTAAGTTGGGTAACGCCA
GGGTTTTCCCAGTCACGACGTTGTAAAACGACGGCCAGTGAATTGTAATACGACTC
ACTATAGGGCGAATTCGAGCTCGGTACCCGGGGATCCTCTAGAGTGGGAGCTCG
GAACACTATCCGACTGGCACCGGCAAGGTCGCTGTTCAATACATGCACAGGATGT
ATATATCTGACACGTGCCTGGAGACTAGGGAGTAATCCCCTTGGCGGTTAAAACG
CGGGGGACAGCGCGTACGTGCGTTTAAGCGGTGCTAGAGCTGTCTACGACCAATT
GAGCGGCCTCGGCACCGGGATTCTCCAGGGCGGCCGCGTATAGGGTCCATCACA
TAAGGGATGAACTCGGTGTGAAGAATCATGCTTTCCTTGGTCATTAGGATCCCGGA
CCTGCAGGCATGCAAGCTTGAGTATTCTATAGTGTCACCTAAATAGCTTGGCGTAA
TCATGGTCATAGCTGTTTCCTGTGTGAAATTGTTATCCGCTCACAATTCCACACAAC
ATACGAGCCGGAAGCATAAAGTGTAAAGCCTGGGGTGCCTAATGAGTGAGCTAAC
TCACATTAATTGCGTTGCGCTCACTGCCCGCTTTCCAGTCGGGAAACCTGTCGTGC
CAGCTGCATTAATG

**Weak side oligonucleotides**
0-linker primer (+/- 5’-Cy3 label)
5’-ACAGGATGTATATATCTGACACGTGCC

77-linker primer
5’-GTACCCGGGGATCCTCTAGAGTG

**Strong side oligonucleotides**
0-linker primer (+/- 5’-Cy3 label)
5’-CTGGAGAATCCCGGTGCCGA

77-linker primer
5’-GATCCTAATGACCAAGGAAAGCATGATTC
